# From treadmill to trails: predicting performance of runners

**DOI:** 10.1101/2021.04.03.438339

**Authors:** B. Crowell

## Abstract

Previous laboratory studies have measured the energetic costs to humans of running at uphill and downhill slopes on a treadmill. This work investigates the extension of those results to the prediction of relative performance of athletes running on flat, hilly, or very mountainous outdoor courses. Publicly available race results in the Los Angeles area provided a set of 109,000 times, with 2200 runners participating in more than one race, so that their times could be compared under different conditions. I compare with the results of a traditional model in which the only parameters considered are total distance and elevation gain. Both the treadmill-based model and the gain-based model have some shortcomings, leading to the creation of a hybrid model that combines the best features of each.

**Author summary:** Running a race on a road allows absolute measures of performance. Trail running, however, has traditionally been thought of as a sport in which the only valid comparison is between different runners competing on the same course on the same day. Even the exact measurement of distance is considered to be unimportant, since courses and conditions vary so much.

An extreme example is the relatively new genre of “vertical” races, in which runners race up a mountain. In a typical example, the competitors cover a horizontal distance of 5 km, while climbing about 1000 m. The winner in one such race had a time almost triple that expected for a state-champion high school runner in a 5k road race. Clearly no comparison can be made here without taking into account the amount of climbing.

In noncompetitive contexts, many runners venture onto mountain trails, lightly dressed and with little equipment, so that it becomes important to be able to anticipate whether they will have the endurance needed to be able to safely complete a planned route. Again, this is impossible without some model of the effect of hill climbing.

## 1 Introduction

This paper presents a method for predicting relative performance on trail runs — “relative” meaning that we can predict the time for course A divided by the time for course B.

Traditionally, runners and hikers have described a trail using two numbers, the horizontal distance and the total elevation gain. For example, if the route is an out-and-back voyage consisting of steady climbing to a peak and a return, then the total elevation gain is simply the elevation of the peak minus the elevation of the trailhead. If the elevation profile of the trip consists of multiple clearly defined ascents and descents, then one adds up the ascents. Although this two-parameter description of the route is easy to derive from a paper topographic map, knowledge of the two numbers is not sufficient to make a very useful estimate of the total energy expenditure.

It has been known for a long time among the officials who measure road races that the effect of elevation change has a nonlinear dependence on the grade. The following argument was advocated by R. Baumel. [1] Consider a closed course whose elevation profile is described by some function *y*(*x*). The derivative *y*^*′*^ is the trail’s slope *i*. The total energy expenditure is an integrated effect of the slope, of the form 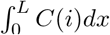, where *C* is a function that describes the energetic cost of running up or down a hill. We will see that *C* has been measured in laboratory experiments, but for the moment we assume only that *C* is a smooth function, so that for small slopes it can be well approximated by the first few terms of its Taylor series, *C*(*i*) *≈ c*_0_ + *c*_1_*i* + *c*_2_*i*^2^. Then for any closed loop over a distance *L*, the contribution from the *c*_1_ term vanishes, and the energy cost is 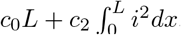. The dependence on the slope is therefore quadratic rather than linear. For example, if we were to exaggerate the elevation profile by a factor of 2, *y→* 2*y*, then the size of the *c*_2_ term would go up by a factor of *four*, not two (in the low-slope limit, on a closed course).

From conversations with runners and hikers, I have found that the result of Baumel’s argument almost always elicits total disbelief, especially when presented as a numerical example showing the extreme smallness of the slope effect when the slope is small. One of the goals of this paper is to test this empirically. As an alternative hypothesis, it is commonly believed that one can get a good measure of the relative energy cost by taking the horizontal distance and adding in a term proportional to the total elevation gain. If the total gain is determined down to a fine enough scale (which with modern technology has become more practical), then this hypothesis is equivalent to the assumption that the cost of running is given by a function of the form

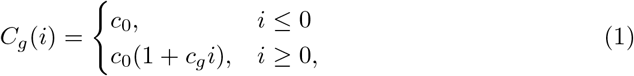

whose graph is shaped like a hockey stick (dashed line in Fig 1). Popularly proposed rules are that 100 m of elevation gain is equivalent to either 400 m or 800 m of horizontal distance, so that *c*_*g*_ is said to be approximately in the range from 4 to 8. There is nothing mathematically impossible about this hypothesis. A function *C*(*i*) of this form evades Baumel’s argument because its hockey-stick shape is not smooth at *i* = 0, and therefore cannot be approximated by its Taylor series.

**Fig 1.**
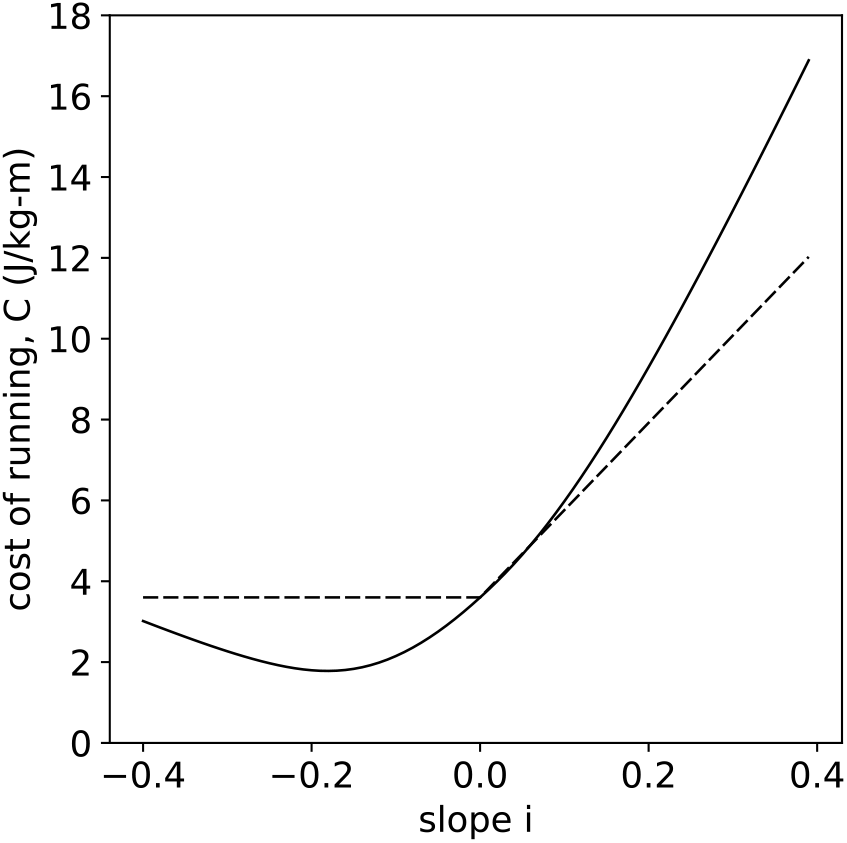
The cost of running as a function of slope. Solid line: the function *C*_*t*_, fit to Minetti’s treadmill data, Eq 5. Dashed line: the function *C*_*g*_, Eq 1, with *c*_0_ chosen to agree with Minetti’s *C*(0) and *c*_*g*_ = 6.0.

In a more sophisticated approach, Minetti *et al*. [8] have used oxygen consumption to measure the energy expenditure of runners on a treadmill at slope *i*, for both running and walking. The results are expressed as *C* = (1*/m*)*dE/ds*, where *m* is the person’s body mass, *E* is the energy expended, and *ds* is the increment of three-dimensional distance, which usually differs negligibly from the increment of horizontal distance *dℓ. C* has units of J*/*kg · m. The correctness of the factor of 1*/m* has empirical support. [10]

Efficiency varies by ∼25% even among elite athletes, [8] [7] and differences are also to be expected between elite and recreational athletes. This is one of the reasons why this study presents a comparative technique, rather than an absolute method for determining a particular runner’s actual energy expenditure in units of kilocalories.

The function *C*(*i*), shown as the solid line in Fig 1, resembles a hyperbola, with a minimum occurring at *i ≈*−0.1 to −0.2. The asymptotes at large positive and negative values of *i* are interpreted in [8] as being determined by the efficiency of eccentric and concentric muscle contraction. For the purposes of this work, a new analytic approximation to the curve found by ref. [8] is used (Appendix 1), and is referred to as *C*_*t*_, where “t” stands for “treadmill.”

Nearly all real-world walking and running is done at −0.2 ≲ *i* ≲ 0.2, where the graph of *C*(*i*) is nearly parabolic.

## 2 Methods

To test these models, I use publicly available race results from the Los Angeles area. This area has a large population and tall mountains. The large population makes it possible to pick out a significant number of runners who have competed in several different races. If the ratio of the runner’s time on courses 1 and 2 is *t*_2_*/t*_1_, then we take this as a measure of the ratio *E*_2_*/E*_1_ of the energy expenditure, which can be compared with the model. It was possible to find courses with a variety of elevation profiles, allowing a test of the dependence of the predictions on the amount of hill climbing. The proportionality of time to energy is found to be good for level and uphill running, but less valid for downhill running, [11] and indeed we will see that these observations seem to hold for the data investigated here.

Table 1 lists the races used as sources of data. One-letter mnemonics are defined so that courses can be referred to succinctly in the text. Because a runner’s performance can change over time due to training and aging, the time period of the study was restricted as much as possible to January 2017 through March 2020 (before the COVID epidemic ended races other than virtual ones in California). Distance and elevation data were analyzed as described in Appendix 3.

**Table 1.**
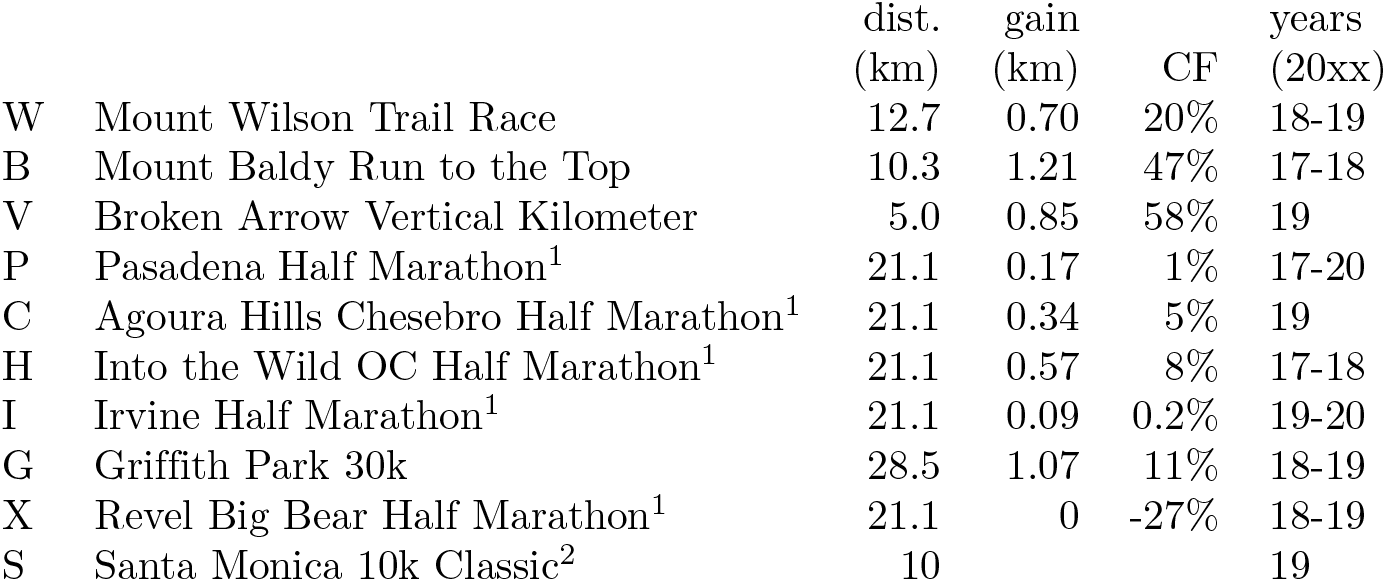
Courses used in this study. Notes: 1. Times up to 2:30 were used. 2. Times up to 1:11 were used for course S. A map of course S was not available, so a flat elevation profile was assumed.

|  |  | dist.<br>(km) | gain<br>(km) | CF | years<br>(20xx) |
| --- | --- | --- | --- | --- | --- |
| W | Mount Wilson Trail Race | 12.7 | 0.70 | 20% | 18-19 |
| B | Mount Baldy Run to the Top | 10.3 | 1.21 | 47% | 17-18 |
| V | Broken Arrow Vertical Kilometer | 5.0 | 0.85 | 58% | 19 |
| P | Pasadena Half Marathon<sup>1</sup> | 21.1 | 0.17 | 1% | 17-20 |
| C | Agoura Hills Chesebro Half Marathon<sup>1</sup> | 21.1 | 0.34 | 5% | 19 |
| H | Into the Wild OC Half Marathon<sup>1</sup> | 21.1 | 0.57 | 8% | 17-18 |
| I | Irvine Half Marathon<sup>1</sup> | 21.1 | 0.09 | 0.2% | 19-20 |
| G | Griffith Park 30k | 28.5 | 1.07 | 11% | 18-19 |
| X | Revel Big Bear Half Marathon<sup>1</sup> | 21.1 | 0 | -27% | 18-19 |
| S | Santa Monica 10k Classic<sup>2</sup> | 10 |  |  | 19 |

Runners’ names and times were obtained by web-scraping public race results, and runners were assumed to be the same person if their first and last names matched. When a runner ran the same race more than once, their best time was used. To avoid biases in comparisons of times in different races, it is necessary here to define an upper limit on the times that will be used from a given race, and to do so in some consistent and unbiased way. Some such limit is in any case defined by race organizers, but is different for different races and usually quite long, often about 4-5 hours for a half-marathon. Competitors who clock the longer times are generally either walking the entire race or alternating between walking and jogging, and especially in more casual races may be pushing a stroller, running alongside their tween-age child, or staying in a costumed group for fun and emotional support. Because the physiological data and models used in this work are not applicable to walking, I impose a somewhat arbitrary time limit of 2.5 hours on half-marathon times. These limits, as well as others, where imposed, are described in the notes in Table 1. For course S, the time limit was derived by scaling down the half-marathon time limit in proportion to the distance. The other courses in this study are of a qualitatively different character, so for them I simply used the race organizers’ cut-off. The resulting bias is an inherent limitation of this work.

Exertion depends most strongly on distance, and the goal of this work is to tease out effects from other factors, which are often weaker. For this reason, distance is a confounding variable in this study and has been controlled for as much as possible by using races at a consistent distance, the half marathon (21.1 km), or distances that, taking extreme climbing into account, result in similar times. These are the distances at which the largest sample sizes are available for mountain trail races. Appendix 5 describes how the remaining inevitable variations in distance have been taken into account, as much as possible.

In table 1, two measures of hilliness are given. The total elevation gain is the only parameter needed in order to calculate an energy expenditure using the function *C*_*g*_. The next column gives a statistic I will refer to as the climb factor, CF, which is defined as the fraction of the runner’s total energy expenditure that is devoted to climbing. That is, if *E* is the actual energy required for the course, and *E*_0_ the energy that would have been required if the race had been perfectly flat, then

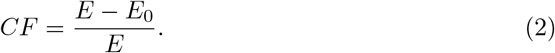

Inverting this equation gives *E* = *E*_0_*/*(1 −*CF*), so that if the horizontal distance is known, a measure of effort can be found by dividing the distance by 1 −*CF* .

To define quantitative tests of the models, consider a comparison of courses 1 and 2. The observed data are the runner’s times *t*_1_ and *t*_2_, and the model predicts the ratio of the energy consumption *E*_1_*/E*_2_. Define

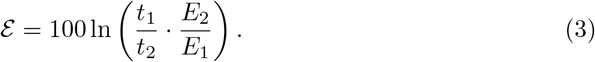

For small errors, *ℰ* is approximately the relative error in the prediction, expressed as a percentage. The use of the logarithm transforms multiplicative sources of error into additive quantities.

We pick a feature of the model that is to be tested. For example, we would like to see whether the model does a good job of predicting the relative times for flat races compared to steep uphill-only races (Fig 2, c). For this example, we make a list of courses that are relatively flat (P, C, H, and I), and a list of some that are steep uphill-only courses (B and V). We then find every case where the same runner did a run *j* from the first list and a run *k* from the second, and compute the error *ℰ*_*jk*_, which will be positive if the runner’s time in the uphill race *k* is overpredicted by the model relative to their time in the flat race *j*.

**Fig 2.**
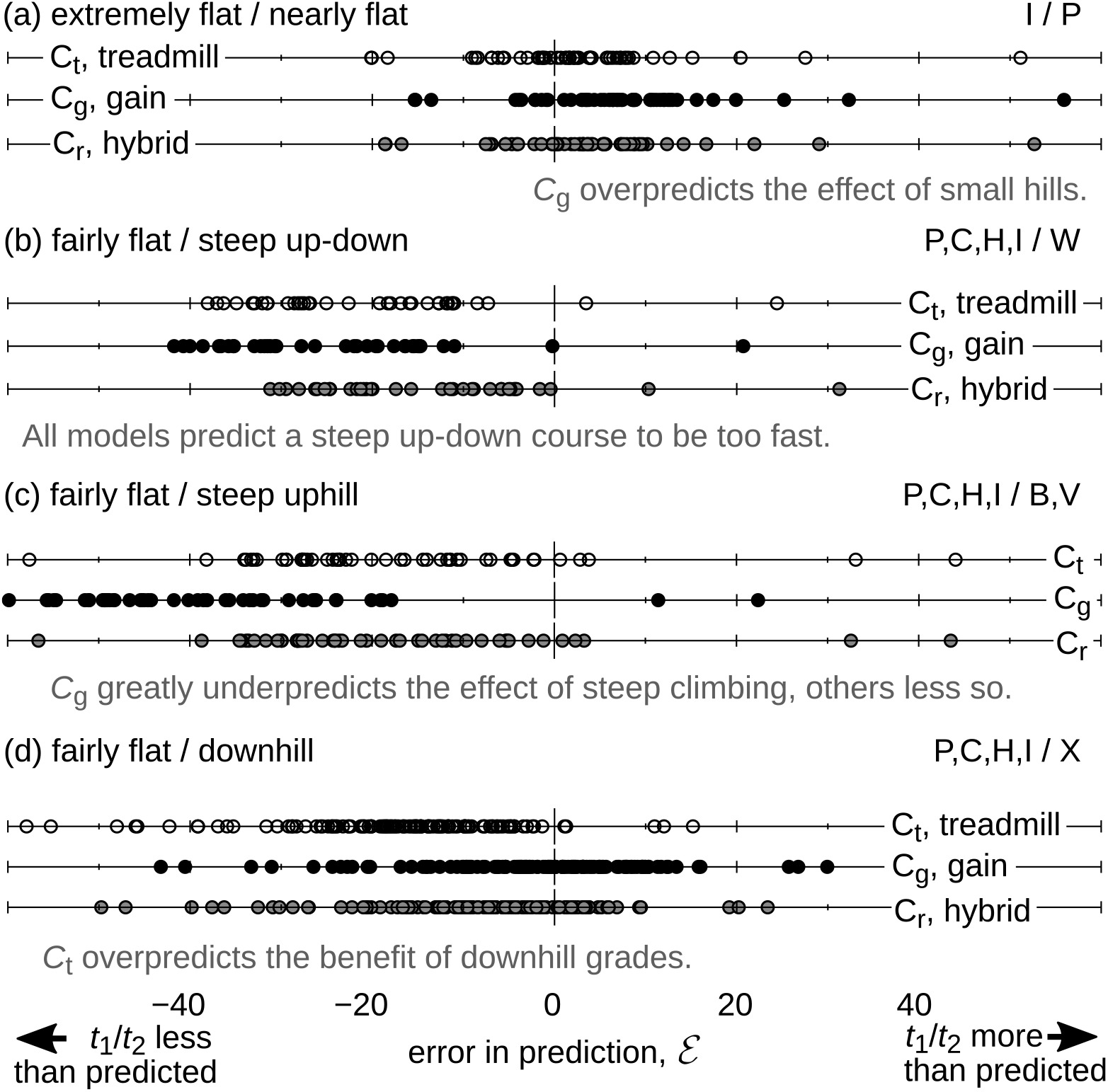
Tests of the predictions of the functions *C*_*t*_ derived from treadmill data (open circles), *C*_*g*_ based on elevation gain (black circles), and the hybrid “recreational” model *C*_*r*_ (gray circles). Positive *ℰ* means that the runner’s time in the first-listed race is greater in reality than in the model.

## 3 Results

Fig 2 shows a comparison of the quality of the predictions of the functions *C*_*g*_ and *C*_*t*_ as descriptions of the effects of going up and down hills. The third model *C*_*r*_ is a hybrid of these, introduced in section 4.2. All predictions were corrected for distance as described in section 5. Each of the four sub-figures a through d has been constructed so as to test a particular feature of these models. We discuss each in turn.

a. Here we compare the extremely flat half-marathon I, having only 90 m of elevation gain, to half-marathon P, which is slightly more hilly with 170 m of gain, or about twice as much. According to the treadmill-based model *C*_*t*_, the effects of climbing and descending nearly cancel out, giving a negligible *CF <* 1% for each run, as expected from Baumel’s argument. In the gain-based model *C*_*g*_, however, the effect of the hills on course P is 6 times its elevation gain, which is equivalent to adding 1.0 km to its length. The effect for I would be half as much, causing the model to predict a considerable difference in the times on the two courses. In the figure we see that Baumel’s approximation is a good one here. The median error for *C*_*t*_ (open circles) is only 1.7%, while that for *C*_*g*_ (filled circles) is +6.4%, the positive sign showing that the effect of the small hills is over-predicted. Of the four tests a-d, this is the only one where the effect being probed is small enough to require statistical analysis rather than simple visual inspection. Such an analysis (Appendix 4) show that systematic error in *C*_*g*_ is significant (*p* = 3 *×* 10^−6^), while any such evidence against *C*_*t*_ is statistically marginal.
b. In this portion of Fig 2, we compare times in a set of fairly flat half-marathons (with CF values ranging from 0.2% to 8%) with course W, a trail race up and down half of Mount Wilson, *CF* = 20%. Although the distance of W is much shorter, most runners’ times are only slightly lower. We see that both models greatly underpredict the runners’ times in the mountain race. Most of the running in this race is on slopes with |*i*| *≈* 0.10 to 0.15. A likely interpretation is that on the uphills, *C*_*g*_ is an underestimate (see c, below), while on the downhills *C*_*t*_ is an underestimate. The race is run on a trail that is mostly a narrow single track, with steep hillsides on the climber’s right. Safety is likely to inhibit many runners from going downhill at anything like the pace that would be possible for the elite mountain runners in ref. [8] on a treadmill, and trail etiquette dictates that they yield the right of way when encountering people who are still on their way up.
c. This test compares runners’ times on the same flattish half-marathons with their performances in two races, B and V, in which runners go up a mountain and finish at the top. Of the sample size of *n* = 32, only one person was on course V, a “vertical kilometer”-style race which was run in Northern California. Although both models systematically underestimated the difficulty of the uphill races (*E <* 0), the underestimate is far more severe for *C*_*g*_ than for *C*_*t*_. Course B consists almost entirely
d. of climbing on grades 0.05 *< i <* 0.25, at which *C*_*g*_ is less than *C*_*t*_ and is apparently a considerable underestimate. Additional factors leading to *ε<* 0 in both cases are certain to include the aerobic challenge of finishing the race at an elevation of over 3000 m, as well as the difficult footing on the final section.
e. Here we compare the same set of fairly flat half-marathons with a road half-marathon, course X, which consists entirely of running *down* a mountain. The downhill race is run along with a marathon, which the race’s organizers advertise as being extremely fast and a good way to achieve a “BQ” or qualifying time for the Boston Marathon. Surprisingly, most runners’ times in the downhill half-marathon were only about 10 minutes shorter than in the flat ones, and quite a few runners in the sample actually took longer for the downhill race. We have already seen evidence in part b above that the high physiological efficiencies observed in the lab for downhill running may not translate proportionately into speed. In W, the issues may have been safety and courtesy, which would not have been relevant in this race on a wide asphalt road. A more likely explanation here is that for recreational runners who have not trained extensively on steep hills, a long downhill of this length can be physically difficult due to the strong eccentric strain on the quadriceps, as well as testing the tensor fascia latae.

## 4 Discussion

### 4.1 Interpretation of results

Fig 3 presents a graphical summary of the interpretation of these results.

**Fig 3.**
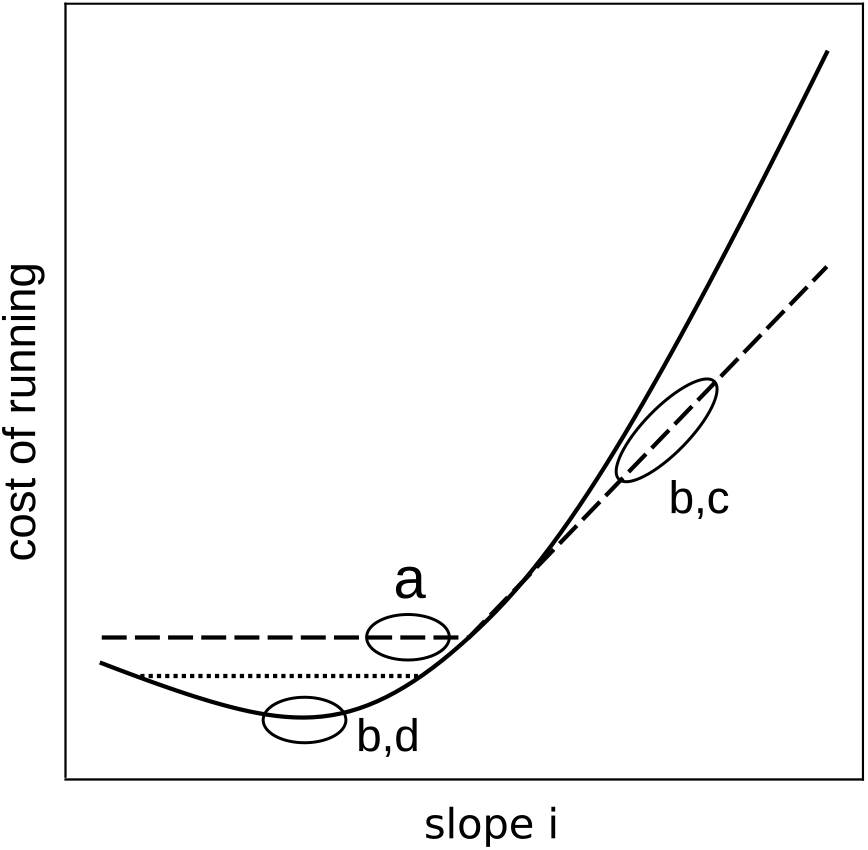
A graphical summary of the interpretation of the systematic errors in the models *C*_*t*_ and *C*_*g*_, observed in parts a, b, c, and d of Fig 2. Portions of the model that are interpreted as being inaccurate are circled and labeled with the test that provides the evidence for the inaccuracy. The interpretation is more conjectural for b, since both models had similar errors, but hypothetically for different reasons. The solid line is the treadmill-based function *C*_*t*_ and the dashed one *C*_*g*_. The dotted line is a modified version *C*_*r*_ of the treadmill function, defined in Eq 4 and found here empirically to be more appropriate for recreational runners in real-world trail conditions.

The observations mainly support the model *C*_*t*_, except that as downhill grades get steeper and steeper, it appears that the speeds of most recreational runners in real-world conditions reach a point of diminishing returns far earlier than would have been imagined from energy measurements in treadmill studies. The results for course X, which has an average *i* ≈ − 0.05, suggest that this point of diminishing returns is reached at relatively small negative slopes, perhaps *i* ≈ − 0.03. There is evidence that humans are unable to run at maximum aerobic capacity on downhill grades, [11] so that we should in fact expect the proportionality of time to energy to break down. But even if this were not so, running fast downhill in real-world conditions is difficult not just because of physiology but also, and probably more importantly, because of safety, etiquette, and training of the quadriceps under eccentric loads. These latter effects cannot be quantified in any universal way. However, it would be irresponsible to provide runners, especially recreational athletes, with scientific advice that would give an unrealistically rosy picture of the difficulty of a run.

### 4.2 Hybrid model

I have therefore investigated some possible modifications to the function *C*_*t*_. A modification that did not work well was to adjust the values of the parameters *p* and *d* given in Table 2 so as to shift the minimum of the function up and to the right. This was unsuccessful, because the smooth analytic character of Eq (5) makes it impossible, by varying its parameters, to dramatically modify the function’s behavior for −0.06 ≲ *i* ≲ −0.03 while retaining its apparently correct behavior at −0.03 ≲ *i* ≲ 0. A more successful ad hoc recipe was simply to introduce a cut-off in *C*, i.e., to define a “recreational” version of the function,

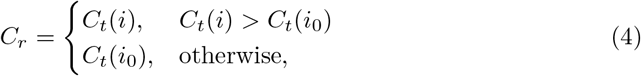

where *i*_0_ = −0.03. In other words, we simply chop the bottom off of the curve of *C*_*t*_, at the dotted line in Fig 3.

**Table 2.** Parameters for Eq (5). These parameters were found by constraining Eq 5 to agree with the polynomial fits in ref. [8] on the following degrees of freedom: the function is minimized at the same *i*, and has the same value of *C* there; the functions agree at *i* = 0. Furthermore, the slopes at *±*∞ were constrained to have the asymptotic values found in that work.

|  | *running* | *walking* |
| --- | --- | --- |
| $a$ | 26.07 J/kg · m | 22.91 J/kg · m |
| $b$ | 0.03104 | 0.02621 |
| $c$ | 1.381 | 1.315 |
| $d$ | -0.06547 | -0.08317 |
| $p$ | 2.181 | 2.209 |

The results of the hybrid model *C*_*r*_ are shown as gray circles in Fig 2 and are in general fairly good. The main remaining inaccuracy is the underprediction of the effect of steep hills in test c. It would be tempting to modify the function *C* to give it an even more severe upward curve for *i* ≳ 0.3. This does not seem warranted by the present data, since other factors may be at work, including altitude and rough footing.

## 5 Conclusions

This paper presents a model that can be used to predict the time of a runner on a course, given their time on some other course. This model is a hybrid of two others that have been previously proposed. Testing against a sample of times clocked by mostly recreational athletes shows that the model usually gives predict corrections to within about 10-20%, even when the route includes extreme climbing or descents. It would be desirable for future work to refine this work by controlling better for factors such as altitude, training, and the quality of footing on mountain trails.

## Appendix 1. Analytic approximation to the treadmill function *C*_*t*_

It is convenient to describe the function *C*(*i*) using a fit to the form

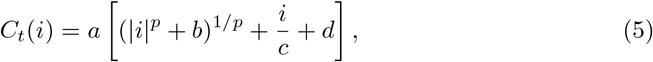

where the subscript *t* stands for treadmill. Parameters fitted to the results of ref. [8] are given in Table 2. The purpose of using this form, rather than the polynomial fit given by [8], is to make the computations degrade gracefully in cases where the limitations of GPS tracks or data from digital elevation models produce unrealistically steep slopes. In such cases, this expression approaches the physiologically expected asymptotic behavior. Although the present work focuses only on running, parameters for walking are presented as well. The results for running are empirically found to be nearly independent of speed, whereas the ones for walking are not. For walking, ref. [8] measured the energy consumption at the speed that was found to be most efficient for that particular subject.

## Appendix 2. Analytic approximation to world-record speeds

Cameron [2] has given a convenient closed-form approximation to world-record speeds of runners at various distances,

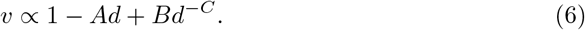

This is shown as the red curve in figure 4. The parameters are given in Table 3

**Table 3.**
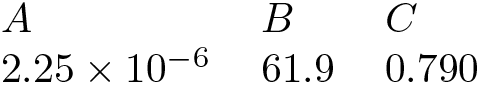
Parameters for Eq (6), for *d* in meters.

**Fig 4.**
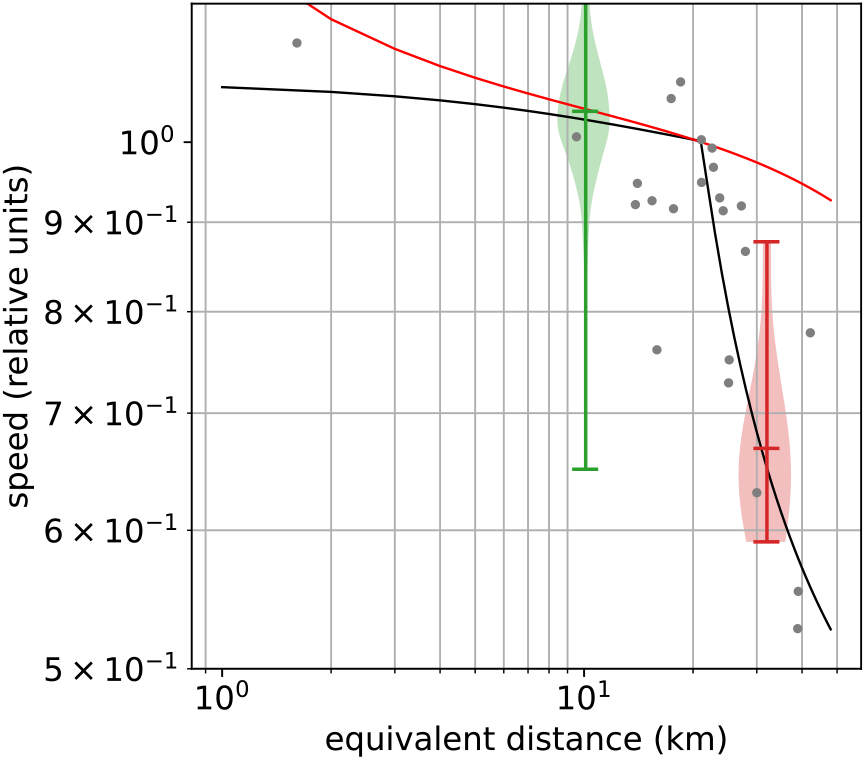
Relative speed versus equivalent distance *d*. All speeds are normalized relative to the speed at half-marathon distance. The black curve is the function defined by Eq 11, with *d*_*c*_ set to a half-marathon distance. The red curve is a fit to world-record times. [2] The green and red violin plots show the distribution of speeds in races S and G relative to the same runners in half-marathon race P (sample sizes 1303 and 11, respectively). The gray dots are the author’s personal-record times from a variety of courses. The equivalent distances were determined from the horizontal distances using the curvilinear function *C*_*t*_(*i*) in Eq 5, which is based on treadmill data.

## Appendix 3: Analysis of elevation data

Digital maps projected into a horizontal plane were obtained from the race organizers’ web site or in some cases by tracing roads and trails in a Google Maps application. Elevation data were obtained from publicly available digital elevation models (SRTM1) having a horizontal resolution Δ*x* = 30 meters. (Elevation data from handheld GPS/GNSS units are more difficult to obtain from public sources, and are in any case of questionable reliability for this purpose, since the uncertainty can be very large when all satellites are near the horizon or when the terrain is rough, causing radio echoes from the walls of canyons.)

The use of these data is inherently subject to certain errors, which need to be minimized. Trails and roads are intentionally constructed so as not to go up and down steep hills, but the DEM may not accurately reflect this. The most common situation seems to be one in which a trail or road takes a detour into a narrow gully in order to maintain a steady grade. If the gully is narrower than the horizontal resolution of the DEM, then the DEM doesn’t know about the the gully, and the detour appears to be a steep excursion up and then back down the prevailing slope.

Empirically, I have found that sensitivity to these effects can be minimized if the elevation profile of the run *y*(*x*) is filtered by convolving it with a rectangular windowing function having width *w* = 200 meters. This tends to eliminate unrealistic glitches in the elevation data, and also seems to give a fairly close reproduction of race organizers’ estimates of total elevation gain. This choice of *w* gives sane results for routes in mountainous terrain, and is used throughout this work, even for flat courses on city streets. For a course that is relatively flat and has many small, short hills, *w* ≈ 60 m gives more accurate results, but I have used the larger value of *w* throughout this work in an effort to maintain consistency.

DEM data are packaged by their providers in rasterized form, much like a digital photo in which each pixel’s “brightness” is the elevation. This models the continuous landscape *z*(*x, y*) as something like a chess board in which each flat square has a different height, with vertical steps occurring along their edges. When a course lies at an angle across such a modeled landscape, the resulting elevation profile *z*(*ℓ*) becomes an irregular staircase. Even if the staircase structure is completely undetectable by eye on a graph of *z* versus *ℓ*, there can be a significant positive systematic error in the energy estimates. The extent to which this error is mitigated by the filtering depends strongly on the ratio Δ*x/w*. Therefore in this work I have used bilinear interpolation to reconstruct a continuous function *z*(*x, y*) from the raster data. Turning the interpolation on and off shows that when Δ*x/w* = 0.15, the interpolation is hardly necessary, causing an additive change *CF*_*o*_ −*CF*_interp_ of about 0.001 to 0.002 on two sample courses, or |Δ*CF/CF*_interp_| ≲ 0.05. However, with Δ*x/w* = 0.5, the use of bilinear interpolation becomes crucial to obtaining good results, with the additive errors rising to 0.01 to 0.03 and relative errors to as big as 0.2 on a flattish course.

The mileage derived from a GPS track can vary quite a bit depending on the resolution of the GPS data. Higher resolution increases the mileage, because small wiggles get counted in. This has a big effect on the energy calculation, because the energy is mostly sensitive to mileage, not gain. For races that were advertised as 5k or half-marathon races, I have therefore used the advertised distance, as shown in Table 1, in order to calculate the first-order estimate of the energy, but have used the elevation gain and CF value derived from the actual GNSS data.

## Appendix 4: Statistical analysis

In section 3, test (a) probes an effect small enough that visual inspection of the scatter plots is not a satisfactory way of testing hypotheses. Specifically, we want to know whether the apparent systematic error in the model *C*_*g*_ is statistically consistent with zero.

We do not know a priori the underlying probaility distribution of the ratio of times or of its logarithm *ℰ*. One might have expected based on previous work [5] that the times would be log-normal, in which case *ℰ*. would be normally distributed. However, a Q-Q plot shows that this is not the case for the present data-set, and in fact the distribution of *E* is asymmetric. The ratio of times, however, has a symmetric and leptokurtic distribution. Its symmetry allows the use of the one-sample Wilcoxon test. For *C*_*g*_ the null hypothesis is rejected with *p* = 4 *×* 10^−6^, while for *C*_*t*_, *p* = 0.07. Thus the defect in *C*_*g*_ is significant, while any such evidence against *C*_*t*_ is statistically marginal.

## Appendix 5: Model of endurance

Animals run more slowly at long distances, and mathematical modeling of this fact dates back about a century. [6] Recent workers have described methods for fitting parameters to the data for individual runners, [9] [3] which for example allows a first-time marathon runner to estimate an appropriate pace. In the present work, there are not enough data available to allow this kind of individualized description of the runners. For this reason, I have concentrated on data from a narrow range of middle distances, with the total energy expenditure being close to that of a flat half-marathon road race. But the endurance required for these races does vary, and this makes it desirable to have some rough method of compensating for the variation in pace with distance. Here I describe a very simple model that has the following characteristics that make it suitable for this study: (1) its two parameters are universal rather than fits to the characteristics of an individual; (2) its dependence on the parameters is purely multiplicative, i.e., varying the parameters only rescales the axes on a graph of speed versus distance. The model is essentially a simplification of the one constructed by Rapoport, [9] with modifications to suit these purposes.

First we compute an equivalent distance *d*, which is the distance of flat running that would require the same energy expenditure as the actual run. If the runner’s time is *t*, then *v* = *d/t* has dimensions of speed, but is in fact a measure of energy per unit time, or power. We then have

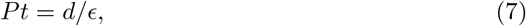

where *P* is the power and *ϵ* is a measure of the runner’s efficiency. For example, a recreational runner with a slight roll of belly fat will have a lower value of *ϵ* because of the increased energetic cost of transporting the additional body weight. Although it would seem that we are now introducing an individualized parameter *ϵ*, the model is designed so that at the end of the calculation, cancellations occur that allow *κ* to be predicted on a universal basis.

The power *P* depends on aerobic fitness and on the proportions of fat and carbohydrates being burned in aerobic metabolism. Fat burning is slower than carbohydrate burning by a factor *β* ≈ 0.4. [9] If we let *f* be the fraction of energy production from carbohydrates, then

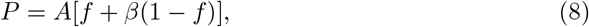

where the proportionality constant *A* is another per-individual parameter that it will be possible to normalize away later. This expression’s linearity in *f* is an approximation to results from real-world data that provide evidence for slightly nonlinear behavior. [9]

The runner’s supply of carbohydrates *c* is limited by the amount of glycogen that can be stored in the liver and the leg muscles. If *f* is chosen optimally, then there will be some distance *d*_*c*_ = *cϵ* that can be run with pure carbohydrate fuel, while longer distances will require *f <* 1. Thus,

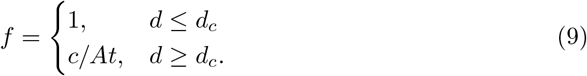

Under these assumptions, the runner’s speed will be the same in races at all distances less than *d*_*c*_, which is unrealistic. We will first work out the consequences of Eq 7-9 and the introduce a simple elaboration that more realistically reproduces the effects of fatigue.

Solving Eq 7-9 and expressing *κ* as a correction factor relative to the short-distance maximum speed *v*_*m*_ = *Aϵ*, we find

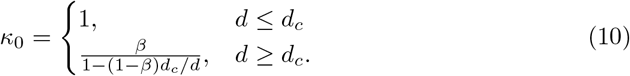

This depends on the universal parameter *β* ≈ 0.4 and also on the critical distance *d*_*c*_. The latter is a measure of endurance and does depend on individual factors such as body composition and training, as well as on strategies such as carbohydrate loading. However, for the sample of recreational athletes studied here, I hypothesize that one can fix a universal value of *d*_*c*_ lying somewhere around the half-marathon distance, and find a reasonable description of real-world data.

It is not true in reality that runners can maintain the same pace at any of the distances below *d*_*c*_, for which glycogen suffices. As the distance increases from 5 km to the half-marathon distace of 21 km, one observes a decrease in speed which, as originally observed by Hill, [6] appears linear on a graph of speed versus the logarithm of distance. In the men’s and women’s world-record times, this decrease is about 5%. The graph then shows a knee, like the one described by Eq 10. The more gradual decrease for distances before the knee is generically described as being due to fatigue, which is a complicated and poorly understood phenomenon involving a variety of factors, many of which are mediated by the central nervous system rather than by any change at the chemical or tissue level. As an *ad hoc* correction, we multiply the result of Eq 10 by a factor controlled by a small parameter *Q*:

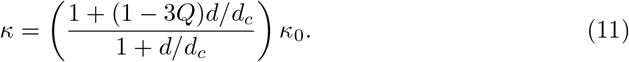

The factor of 3 is introduced so that *Q* is approximately equal to the reduction in speed between a 5k and a half-marathon, and we set *Q* = 0.05.

Empirically, for the mostly recreational runners studied here, a reasonable description of the data is achieved when *d*_*c*_ is set to the half-marathon distance, which is the value adopted in this work. Fig 4 shows that setting *d*_*c*_ to half-marathon distance gives a good fit to some real-world data.

Although the fit in Fig 4 is rather good, much higher values of *d*_*c*_ might be more appropriate for higher-level endurance runners. For example, Eliud Kipchoge’s personal-record speed is only 8% lower in the marathon than in the 1500 m. This can only be reproduced in this model if *d*_*c*_ is roughly marathon distance for him. For high-level running competitions, Gardner and Purdy give a method of comparing with a standard performance curve, based on a compilation of world records. [4] The red curve in Fig 4 is an approximation to the locus of world-record times (Appendix 2).

Although fitting parameters to individual runners’ characteristics is not the main purpose of this work, doing so is very easy with the model 11, due to its purely multiplicative structure. When data are viewed in the format used in Fig 4, as a log-log plot of speed versus distance, the standard curve *κ*(*d*) is simply slid around horizontally and vertically to match the data, which has the effect of determining the runner’s *v*_*m*_ and *d*_*c*_.

## Acknowledgements

The author thanks Wendy Yen for helpful conversations about statistical tests.

## Data availability statement

The compilation of race times was derived from public sources and is itself publicly available at https://github.com/bcrowell/trail. The code used to analyze the data is contained in that repository and in https://github.com/bcrowell/kcals. All code is under a GPL license. I have also used Zenodo to assign a DOI to the data: 10.5281/zenodo.4661554.

## References

1. Baumel R. Hill effect to second order. Measurement News. 1989 Jan;33:36.

2. Cameron DF. Time-equivalence Model. 1998 Jun. Available from: http://www.cs.uml.edu/∼phoffman/cammod.html

3. Emig T and Peltonen J. Human running performance from real-world big data. Nat Commun. 2020 Oct 6;11:4936. doi:10.1038/s41467-020-18737-6

4. Gardner J and Purdy J. Computer generated track scoring tables. Medicine and science in sports. 2, 152–161. 1970;2(3):152.

5. Godsey B. Comparing and forecasting performances in different events of athletics using a probabilistic model. Journal of Quantitative Analysis in Sports. 2012 Jun;8(2):1. doi:10.1515/1559-0410.1434

6. Hill AV. The physiological basis of athletic records. Lancet. 1925;5:481–486.

7. Lucia A, Olivan J, Bravo J, Gonzalez-Freire M, and Foster C. The key to top-level endurance running performance: a unique example. British Journal of Sports Medicine. 2008;42(3):172.

8. Minetti AE, Moia C, Roi GS, Susta D, and Ferretti G. Energy cost of walking and running at extreme uphill and downhill slopes. J Applied Physiology. 2002;93:1039.

9. Rapoport BI. Metabolic factors limiting performance in marathon runners. PLoS Comput Biol. 2010 Oct;6(10):e1000960.

10. Shaw AJ, Ingham SA, Folland JP. The valid measurement of running economy in runners Med Sci Sports Exerc. 2014 Oct;46(10):1968–73.

11. Vernillo G, Giandolini M, Edwards WB, Morin JB, Samozino P, Horvais N, Millet GY. Biomechanics and physiology of uphill and downhill running. Sports Medicine. 2017 Apr;47(4):615–29.

